# Graph-guided flow matching enables biologically informed prediction at single-cell resolution

**DOI:** 10.64898/2026.11.12.766125

**Authors:** Huai-Yu Liou, Maximilian Sprang, David A. Selby, Gerrit Grossmann, Sebastian Vollmer, Pablo Riesgo-Ferreiro, Jennifer V. Gerbracht, Barbara Schrörs, Boris Strilic, Johnny Kim

**Affiliations:** TRON – Translational Oncology at the University Medical Center of the Johannes Gutenberg University gGmbH, Mainz, Germany; Faculty of Biology, Johannes Gutenberg University of Mainz, Mainz, Germany; Department of Dermatology, University Medical Center of the Johannes Gutenberg University Mainz; Institute of Quantitative and Computational Biology, Johannes Gutenberg University, Mainz, Germany; Data Science and its Applications, German Research Center for Artificial Intelligence (DFKI), Germany; BioNTech SE, Mainz, Germany; Vascular Biology and Therapeutics Program and Department of Comparative Medicine, Yale University School of Medicine, New Haven, CT, USA

## Abstract

Perturb-seq enables systematic interrogation of massively parallel genetic perturbations at single-cell resolution, yet exhaustive characterization of the full perturbation space remains impractical. Here, we present Graph-guided Flow Matching (GFM), a generative framework that integrates biological knowledge graphs with latent flow matching to predict post-perturbation single-cell transcriptomes. Across multiple large-scale Perturb-seq datasets, GFM outperforms a strong linear baseline and current deep-learning models in predicting unseen single-gene perturbations, while training up to 30x faster and remaining robust with limited training data. GFM also demonstrates competitive performance in the two-gene perturbation prediction task when neither perturbed gene is observed during training. Ablation experiments show that combining flow matching with knowledge graphs enables accurate prediction of perturbation responses. Finally, GFM accurately calculates transcriptional consequences of HSP90AA1 inactivation and identifies target candidates unseen by the model in chronic myeloid leukemia. Our study establishes GFM as a biologically informed generative framework that predicts single-cell perturbation responses for prioritization of therapeutic targets.

## Introduction

Perturbation sequencing (Perturb-seq) is a recently developed high-throughput screening technology that combines CRISPR-based gene activation or repression with single-cell RNA sequencing (scRNA-seq). This assay enables systematic examination of hundreds of thousands of genetic perturbations and their effects on the transcriptome of individual cells, facilitating the study of gene-gene interactions, cell differentiation processes, and target identification in drug development^1–5^. Although Perturb-seq captures causal relationships between target genes and their downstream effectors, experimentally testing all possible genetic perturbations and their combinations remains impractical because the complexity of the perturbation space rapidly outpaces experimental scalability. Therefore, computational tools capable of learning from existing Perturb-seq data and generalizing to unseen perturbations are instrumental for scalable target discovery and a deeper understanding of complex gene regulatory networks^6^.

Recent perturbation prediction methods leverage pretrained foundation models to improve generalization to unseen perturbations. For example, single-cell generative pre-trained transformer (scGPT)^7^ adapts a transformer pretrained on observational scRNA-seq data, whereas CellFlow^8^ incorporates representations from a protein language model (ESM2^9^). However, these approaches do not explicitly model perturbation-specific gene-gene interactions and therefore have limited capacity to capture the complexity of gene regulatory cascades induced by genetic perturbations. Recently, biological priors have been integrated into perturbation prediction frameworks using graph neural networks (GNNs). GEARS^10^ incorporates Gene Ontology (GO) and gene co-expression network using GNNs to predict responses to unseen genetic perturbations, whereas TxPert^11^ further incorporates protein-protein interaction (PPI) networks, a data-derived perturbation graph, and image-based features. Despite these advances, the contributions of biological priors to predictive performance remain poorly understood, and their incorporation into an effective generative framework is still lacking.

Notwithstanding, advances in generative modeling have provided new opportunities for predicting cellular responses to unseen perturbations. Flow matching is a recently emerged generative modeling framework that learns continuous transformations, parameterized by neural ordinary differential equations (ODEs), from noise or arbitrary source distributions to target data distributions^12,13^, making it well-suited for single-cell data generation. Therefore, flow matching has been increasingly applied to model cellular dynamics and to interpolate or extrapolate to unobserved cellular states in tasks such as trajectory inference^14,15^ and perturbation prediction^8^. However, a central challenge for flow matching is that the trajectories it seeks to learn are never directly observed: single-cell perturbation experiments provide only separate populations of control and perturbed cells and not paired observations of the same cell before and after perturbation. In addition, existing flow matching approaches remain limited in their ability to incorporate biological knowledge and generalize to unseen perturbations. To address this challenge, Meta Flow Matching^16^ employs a GNN to condition the source distribution, whereas MixFlow^17^ learns a condition-dependent Gaussian-mixture source distribution. Despite these advances, these methods primarily improve how the flow is initialized, rather than how perturbation-induced changes are modeled throughout the transformation. Consequently, they do not explicitly model how perturbations propagate through biological systems, limiting their ability to generalize to unseen perturbations.

Here, we present Graph-guided Flow Matching (GFM), a generative framework that combines multiple biological knowledge graphs with latent flow matching to predict single-cell transcriptomic responses to unseen perturbations. GFM employs latent flow matching with a Gaussian source distribution, classifier-free guidance, and condition-specific optimal transport (OT) couplings to efficiently learn perturbation-induced transcriptomic changes. Through comprehensive benchmarking, we demonstrate that GFM achieves superior performance in single-gene perturbation prediction and competitive performance in two-gene perturbation settings. We functionally validate GFM predictions and show that GFM prioritizes therapeutic targets for chronic myeloid leukemia (CML).

## Results

### GFM model overview

GFM is an end-to-end framework that takes scRNA-seq data with perturbation labels during training and generates post-perturbation gene expression profiles at single-cell resolution. Inspired by prompt-guided generation in modern image synthesis models^18–20^, GFM aims to transform Gaussian noise into perturbed cellular states in a learned latent space conditioned on a knowledge-graph-derived perturbation representation.

Specifically, GFM is built on three major components: a variational autoencoder (VAE), a GNN module, and a neural ODE module. The VAE learns a latent representation from log-normalized single-cell gene expression data while preserving the ability to reconstruct gene expression profiles from the latent space (Fig. 1A). We adapted the GO graph from GEARS^10^ but additionally constructed a PPI graph from the STRING database^21^ and a perturbation similarity graph derived from a K562 genome-wide CRISPRi Perturb-seq dataset^3^. These three graphs are combined and parameterized by a graph attention network (GAT), allowing for biologically informed inductive bias. Each gene in the graph is associated with a trainable perturbation embedding that propagates to neighboring genes, essentially enabling the model to generalize to unseen genetic perturbations. Importantly, the perturbation embedding guides the velocity field generated by the neural ODE module that transports Gaussian noise toward the corresponding perturbed cell state, allowing the model to learn perturbation-induced shifts in small steps rather than directly generating data points (Fig. 1A).

**Fig. 1.**
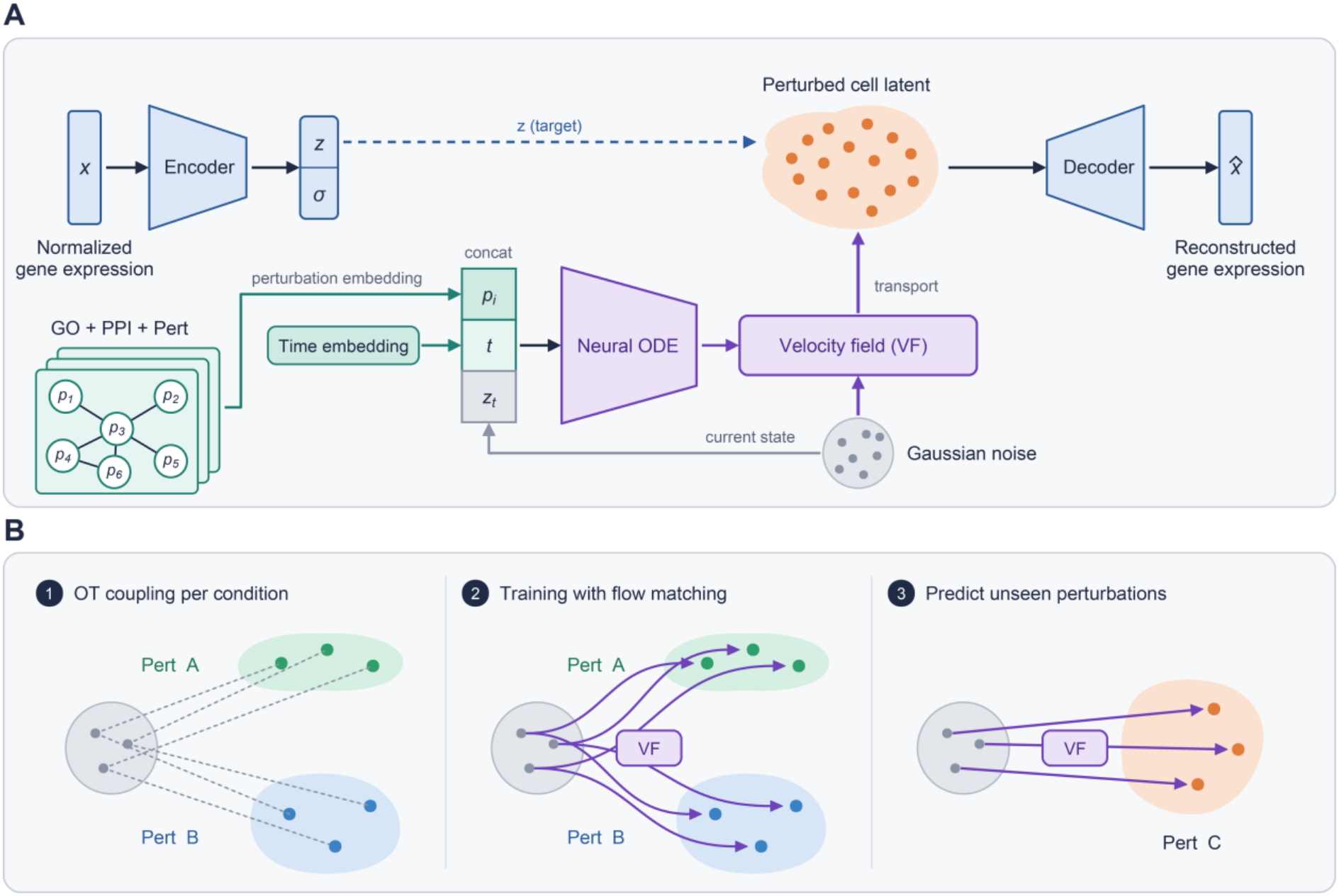
Overview of GFM model architecture and the flow matching training procedure. **(A)** The VAE encodes a log-normalized gene expression profile *x* into the latent code *z*, which can be reconstructed to the gene space as 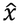 by the decoder. The neural ODE learns to generate a time-dependent velocity field through a predefined path *z*_*t*_ from Gaussian noise (*t* = 0) to a perturbed cell state (*t* = 1), conditioned by a selected GNN-derived perturbation embedding *p*_*i*_ and a time embedding *t*. **(B)** The OT maps are computed independently for each condition before training. Then, the model is trained to generate the velocity field that follows these OT paths. Finally, a trained model can be used to generate unseen perturbed cell profiles. Pert: perturbation.

The VAE module is pretrained independently of the perturbation labels, and the GNN and neural ODE modules are trained jointly using a flow matching objective. Before the flow-matching training, we defined perturbation-specific probability paths between Gaussian noise and perturbed cells as straight interpolants coupled by OT. The neural ODE that is conditioned on the GNN-derived perturbation embeddings is then trained to match the velocity field along these OT paths. Once trained, GFM predicts cellular states under unseen perturbations in the latent space and decodes them back into gene expression space (Fig. 1A). Taken together, GFM is a generative framework for predicting single-cell transcriptomic responses to unseen perturbations by integrating biologically informed conditioning with perturbation-specific OT-based flow matching.

### GFM accurately predicts unseen single-gene perturbation responses

In order to test its generalization to unseen single-gene perturbations, we benchmarked GFM on four CRISPRi Perturb-seq datasets targeting thousands of genes in the chronic myeloid leukemia cell line K562, the retinal pigment epithelial cell line RPE1^4^, the hepatocellular carcinoma cell line HepG2, and the T-cell leukemia cell line Jurkat^5^. For each dataset, perturbations were split into training (75%), validation (10%), and test (15%) sets across three independent data splits (Fig. 2A). GFM was compared with current models, including CellFlow^8^, GEARS^10^, and scGPT^7^. Additionally, a linear baseline model that learns to linearly map a data-driven PCA-derived gene embedding to the perturbation-induced gene expression shift was included^22^ (see Methods). To provide empirical upper- and lower-bound references, we evaluated a technical duplicate (Dup), in which half of the ground-truth perturbed cells served as the prediction for the other half^23^, and a Perturb train baseline that randomly samples perturbed cells from the training set^24^. Model performance was assessed using complementary metrics that capture both mean expression prediction accuracy (Pearson correlation, Pearson delta, and mean squared error (MSE)) and distributional similarity between predicted and observed perturbation responses (2-Wasserstein distance (2-WD) and energy distance (E-distance)). To facilitate comparison of perturbation-specific prediction performance across conditions, all metrics except Pearson delta were normalized by subtracting the corresponding perturb-train baseline values. Pearson delta was exempt from this normalization as it is inherently defined relative to the perturb-train baseline. Unless otherwise stated, evaluation was restricted to the top 20 differentially expressed genes (DEGs) for each perturbation to focus on perturbation-specific transcriptional responses.

**Fig. 2.**
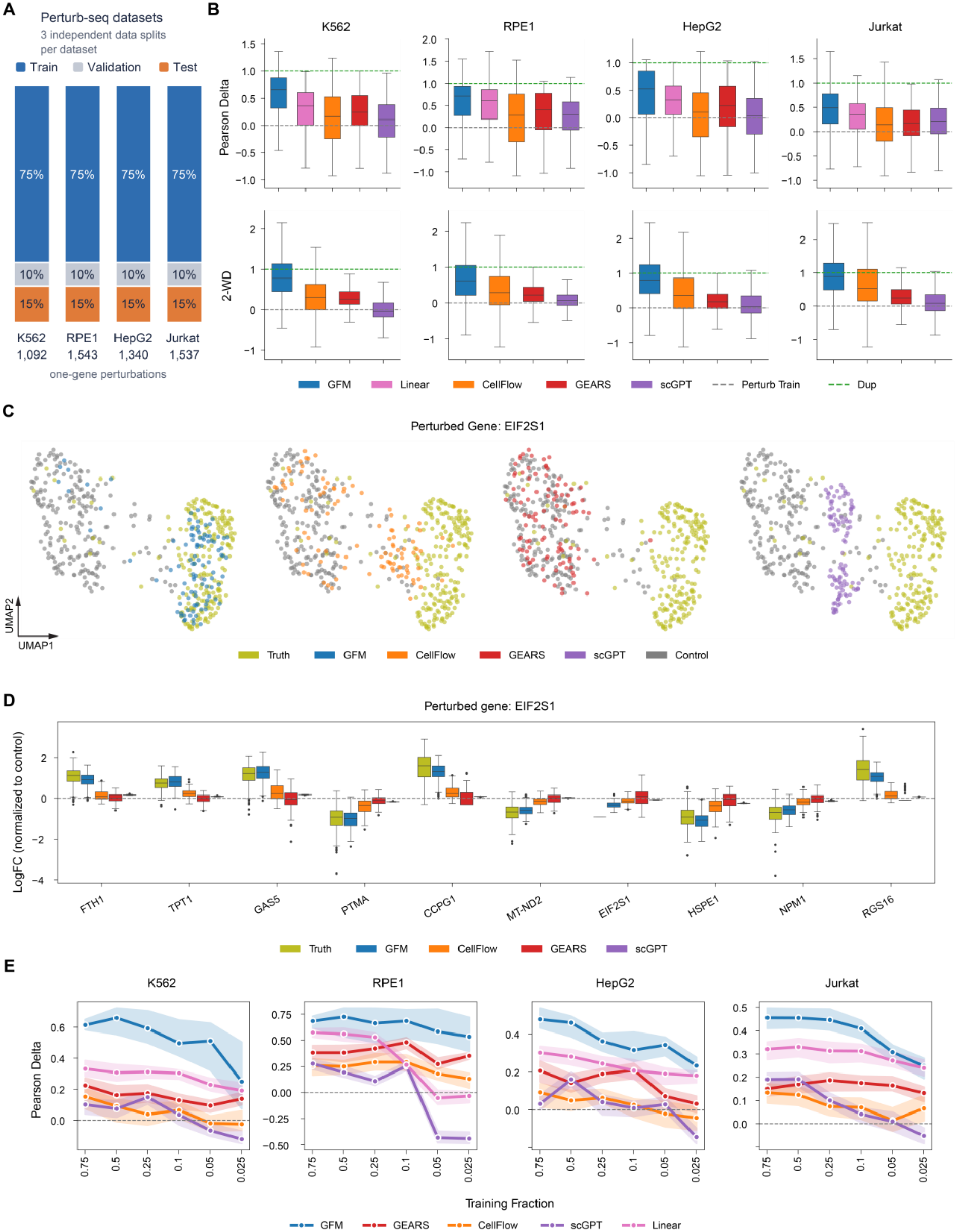
Comprehensive benchmarking of GFM performance in single-gene perturbation prediction. **(A)** The data splits for training and the number of one-gene perturbations in each dataset. **(B)** Box plots showing Pearson delta and 2-WD of the predicted top 20 DEGs of each perturbation for each model in K562, RPE1, HepG2, and Jurkat datasets. Randomly selected perturbed cells from the training set (Perturb train) and technical duplicate (Dup) were used to normalize the metrics. The tested perturbations in three independent data splits were pooled. **(C, D)** Joint UMAP representation **(C)** and the predicted top 10 DEGs **(D)** of EIF2S1 knockdown in K562 from different models. Expression levels were shown as log fold change relative to the mean control cell expression. Truth: ground-truth gene expression. **(E)** Line plots showing Pearson delta of the predicted top 20 DEGs of each perturbation with reduced training fractions (0.75, 0.5, 0.25, 0.1, 0.05, and 0.025). The analysis was performed in the same way for each model in the K562, RPE1, HepG2, and Jurkat datasets. Results calculated from three independent data splits were pooled, and the median and 95% confidence interval were shown.

In all tests, GFM systematically outperformed the other models in predicting the top 20 DEGs at both the mean and population levels. Interestingly, consistent with previous reports^22^, the linear baseline outperformed CellFlow, scGPT, and GEARS for mean gene expression prediction. However, GFM surpassed the linear baseline (Fig. 2B, Fig. S1). Furthermore, GFM consistently yielded lower 2-WD scores (Fig. 2B), indicating that its predicted cell populations more closely resemble true perturbed distributions. However, GFM and other tested models exhibited similar or higher E-distance compared to randomly sampled perturbed cells (Fig. S1), suggesting that none of the models adequately captured the overall dispersion of the perturbed population. Similarly, when performance was assessed using the top 10 to 100 DEGs as evaluation features, GFM demonstrated superior performance across all metrics except E-distance (Fig. S2). We next assessed prediction accuracy by visualizing representative perturbations from the test sets in two-dimensional UMAP space. For example, following EIF2S1 knockdown in K562, GFM-predictions closely overlapped with the observed perturbed cell population, whereas GEARS’ predictions resembled more closely unperturbed cells and predictions based on CellFlow and scGPT failed to overlap with the ground truth (Fig. 2C). Visualization of the predicted top 10 DEGs following EIF2S1 knockdown similarly showed that GFM more accurately captured both the mean and variance of gene expression levels than alternative models (Fig. 2D). Comparable prediction accuracy was observed across perturbations in additional cell lines, including NHP2 knockdown in RPE1, PSMC4 in HepG2, and GRPEL1 in Jurkat ( Figs. S3A-C).

Next, we examined the robustness of GFM under reduced training data. To this end, we evaluated its ability to predict the held-out test set as the training size was gradually reduced from 75% to 2.5%, across three independent data splits (Table S1). As expected, performance declined with a decreasing amount of training data for all models. Notably, GFM consistently outperformed CellFlow, scGPT, GEARS, and the linear baseline across mean metrics and 2-WD, (Fig. 2E, Fig. S4).

Furthermore, GFM required substantially less training time than the other deep-learning models (1.5-30× faster), while achieving inference times comparable to GEARS and ∼100× shorter than those of CellFlow and scGPT (Fig. S5). Taken together, these results establish GFM as a general framework for predicting unseen perturbation-induced transcriptional responses. By integrating a latent flow-matching generative framework with perturbation-specific learning and knowledge graphs, GFM generalizes across perturbations in diverse cell types while consistently outperforming existing methods, including a competitive linear baseline even under limited training data regimes.

### GFM predicts two-gene perturbations

Next, we applied GFM to a CRISPRa Perturb-seq dataset comprising 131 two-gene and 152 one-gene perturbations in K562 cells^3^. Here, we only implemented the GO and PPI graphs in GFM since the perturbation similarity graph was constructed based on a CRISPRi dataset^4^, which intrinsically carries different perturbation-specific information from the CRISPRa dataset. The two-gene perturbations in the held-out test set were further divided into three subgroups based on the number of perturbed genes seen during training: both genes seen (seen-2), one gene seen (seen-1), or neither gene seen (seen-0), reflecting increasing prediction difficulty. For the seen-2 subgroup, we additionally implemented an additive baseline, in which the predicted expression change is defined as the sum of the two corresponding single-gene perturbation changes observed in the training data.

GFM achieved similar performance to the other models in the seen-2 and seen-1 subgroups but showed superior performance in the seen-0 subgroup, where neither perturbed gene was present during training (Fig. 3A, Fig. S6). For example, for the dual activation of CBL and PTPN9, GFM-predicted cells overlapped more closely with the ground truth in 2D UMAP space (Fig. 3B), and the top 10 DEGs predicted by GFM and GEARS were better aligned with the observed data compared to CellFlow and scGPT (Fig. 3C). Consistent with previous reports^22^, the additive model achieved the highest performance for predicting the top 20 DEGs in the seen-2 subgroup, suggesting that the transcriptional effects of two-gene perturbations in this dataset are largely additive. In summary, GFM achieves comparable performance when one or both genes are seen and predicts two-gene perturbations better than current deep-learning models when neither perturbed gene is seen during training.

**Fig. 3.**
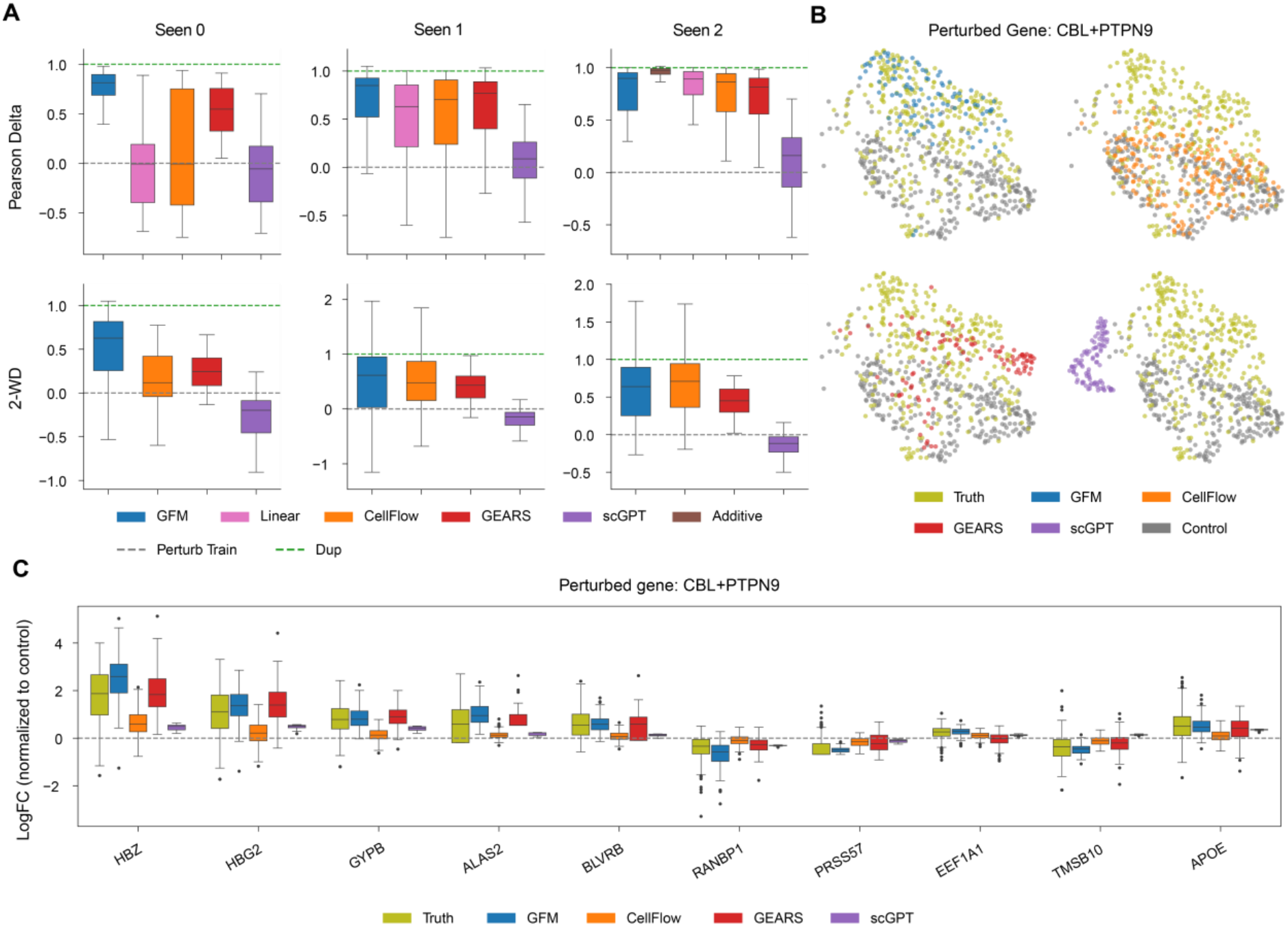
Two-gene perturbation predictions. **(A)** Box plots showing Pearson delta and 2-WD of the predicted top 20 DEGs of each perturbation for each model in seen-0, seen-1, and seen-2 subgroups. Randomly selected perturbed cells from the training set (Perturb train) and technical duplicate (Dup) were used to normalize the metrics. The tested perturbations in three independent data splits were pooled. **(B, C)** Joint UMAP representation **(B)** and the predicted top 10 DEGs **(C)** of CBL and PTPN9 overexpression in K562 from different models. Expression levels were shown as log fold change relative to the mean control cell expression. Truth: ground-truth gene expression.

### Biological knowledge graph strongly influences GFM’s generalizability

To assess how the flow matching framework and the knowledge graph-constrained GNN contribute to GFM’s single-gene perturbation prediction, we performed ablation experiments using the same K562, RPE1, HepG2, and Jurkat Perturb-seq datasets. To ablate the flow-based prior while maintaining the same model structure, we used the same multilayer perceptron (MLP) architecture as in the neural ODE module but took its output directly as the generated latent data points instead of the velocity field. Furthermore, no OT coupling between the Gaussian noise and the target perturbed cells was computed before training. Removing the OT flow matching objective resulted in an overall performance decline, particularly in the distributional metrics (Fig. 4A, Fig. S7). These results highlight the importance of the flow matching generative framework for modeling accurate perturbation effects while preserving single-cell heterogeneity. To measure the contribution of prior biological knowledge, we randomly permuted the knowledge graph while maintaining the number of edges. This manipulation removes biological relationships while preserving comparable graph density and message-passing capacity. Intriguingly, we observed a significant performance loss in both mean and distributional metrics when the graph connectivity became random (Fig. 4A, Fig. S7). Similarly, simultaneous ablation of both components further impaired GFM performance. These data demonstrate that the biological prior knowledge is largely responsible for the generalizability of GFM and that the GNN module and the flow matching component synergistically contribute to its superior predictive performance.

**Fig. 4.**
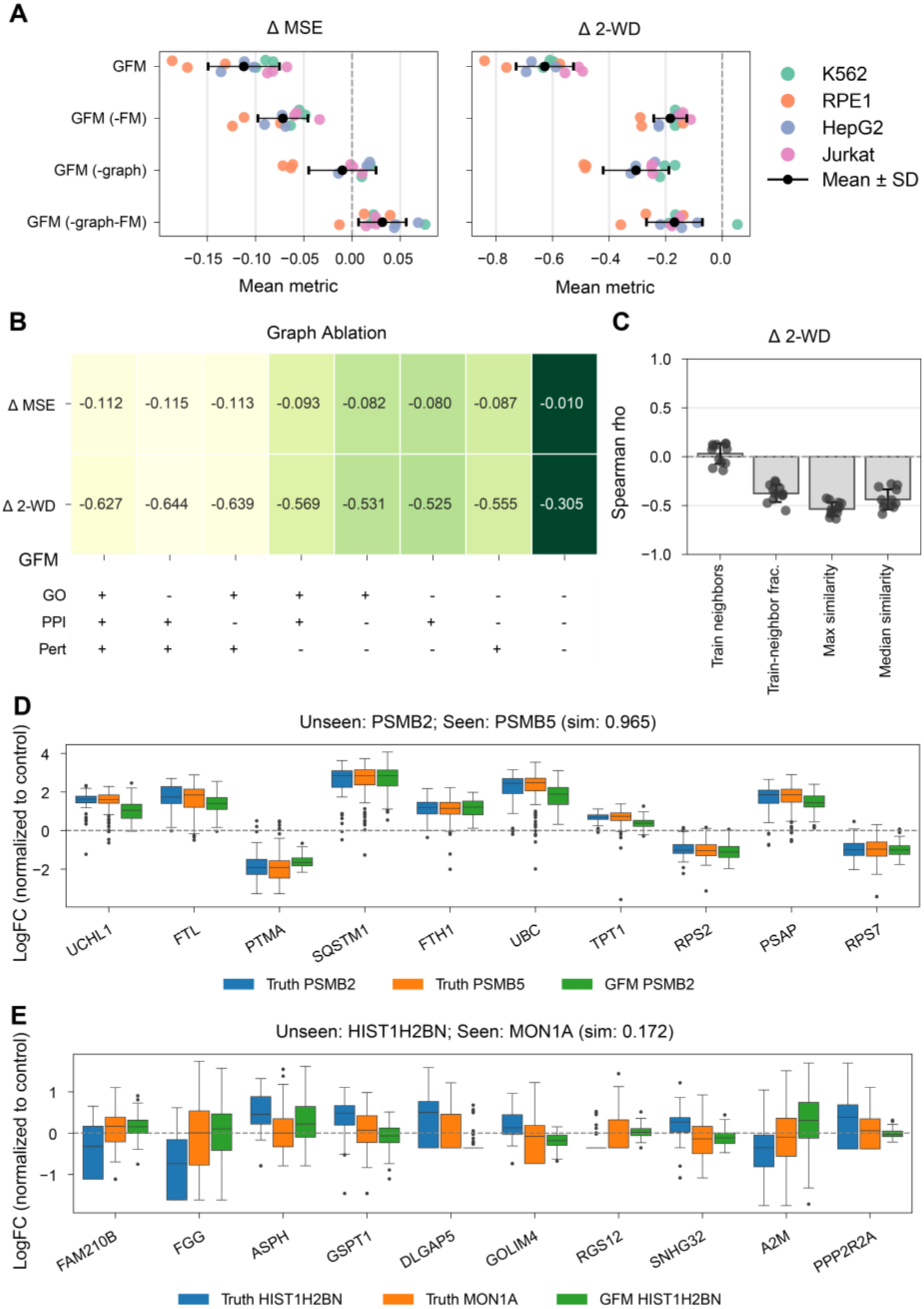
Ablation and graph connectivity analysis for single-gene perturbation prediction. **(A)** The aggregated mean delta MSE and delta 2-WD relative to Perturb train baseline for the three data splits in the K562, RPE1, HepG2, and Jurkat datasets in different ablation conditions. Additional means and standard deviations are shown for each ablation condition. -FM: without OT flow matching, -graph: randomly permuted graph. **(B)** A heatmap showing the averaged delta MSE and 2-WD relative to Perturb train baseline across three independent splits of the K562, RPE1, HepG2, and Jurkat datasets under multiple graph ablation conditions. The indicated biological graph was randomly permuted (-) or remained intact (+). **(C)** Summarized Spearman correlation between delta 2-WD and the graph-derived properties. Each data point shows the Spearman rho of a data split in the K562, RPE1, HepG2, and Jurkat datasets. For each comparison group, the Fisher z-transformed mean of the Spearman rho was shown as the bar, with the error bar showing the standard deviation. **(D, E)** The top 10 DEGs of PSMB2 knockdown in the RPE1 dataset and HIST1H2BN knockdown in the HepG2 dataset, together with the corresponding GFM predictions. The most similar graph-connected perturbations in the training set, PSMB5 and MON1A knockdown, respectively, are also shown. Expression levels are shown as log fold change relative to the mean control cell expression. Truth: ground-truth gene expression. sim: cosine similarity.

We next assessed the contribution of different types of biological knowledge to the predictive performance of GFM by individually ablating each graph component by randomly permuting its connections while leaving all other modules unchanged. Ablation of the perturbation similarity graph markedly impaired GFM performance, whereas ablation of either the GO or PPI graph alone had only modest effects (Fig. 4B, Fig. S8). Simultaneous permutation of any two graphs led to a further decline in predictive performance, indicating that no single graph can compensate for the loss of the others (Fig. 4B, Fig. S8). Together, these results suggest that the PPI and GO graphs provide complementary biological information that is essential for its baseline predictive performance. The perturbation similarity graph, which directly encodes perturbation-specific relationships, appears to be the primary contributor to GFM’s superior generalizability.

To further understand how biological priors influence the predictive performance of GFM, we investigated the correlations between several graph connectivity-derived properties and prediction accuracy. Importantly, since the GAT is a message-passing GNN, a representation of each target node is iteratively shaped by information propagated from neighboring source nodes across layers. Consequently, the quality of a target node representation is expected to depend on its connectivity within the graph^25^. We therefore quantified how the connectivity of a target node, representing an unseen perturbation in the test set, to other perturbation nodes in the graph influences prediction accuracy. To this end, we first measured two intrinsic graph properties for each test perturbation: the number of its neighbors that belonged to the training set and the fraction of all its neighbors that belonged to the training set. We found that the trained-neighbor fraction, rather than the number of trained neighbors, were associated with better generalization performance across test metrics (Fig. 4C, Fig. S9). We then used the ground-truth gene expression data to compute two extrinsic properties between each test perturbation and its training-set neighbors: the maximum and median similarity of their perturbation effects, where perturbation effect was defined as the change in average gene expression relative to control cells and similarity was measured by cosine similarity between these expression-change vectors. Strikingly, these extrinsic properties showed even stronger associations with test performance (Fig. 4C, Fig. S9), indicating that GFM predicts unseen perturbations more accurately when its training-set neighbors have more similar transcriptional effects. For example, the most similar training-set neighbor of PSMB2 knockdown was PSMB5 knockdown (cosine similarity = 0.965), corresponding to an accurate GFM prediction (Fig. 4D). By contrast, the most similar training-set neighbor of HIST1H2BN knockdown was MON1A knockdown (cosine similarity = 0.172), corresponding to poor prediction performance (Fig. 4D). This finding confirms that graph construction that is based on perturbation similarity is more informative for perturbation prediction.

### GFM reproduces HSP90AA1 knockdown effects in CML

Finally, we trained the GFM model on the genome-wide CRISPRi Perturb-seq dataset in K562 cells^4^ and predicted the transcriptional effects of all possible single-gene perturbations. The predicted functions of each perturbation were annotated based on its down-regulated genes, identifying categories such as cytoplasmic translation, RNA processing, and regulation of gene expression (Fig. S10). Focusing on the heat-shock protein HSP90AA1, a well-established therapeutic target that stabilizes the BCR-ABL1 complex and promotes chronic myeloid leukemia progression^26^, GFM correctly predicted the suppression of chaperone-related functions and telomere maintenance following its knockdown^27^ (Fig. 5A) despite HSP90AA1 not being included in the training data.

**Fig. 5.**
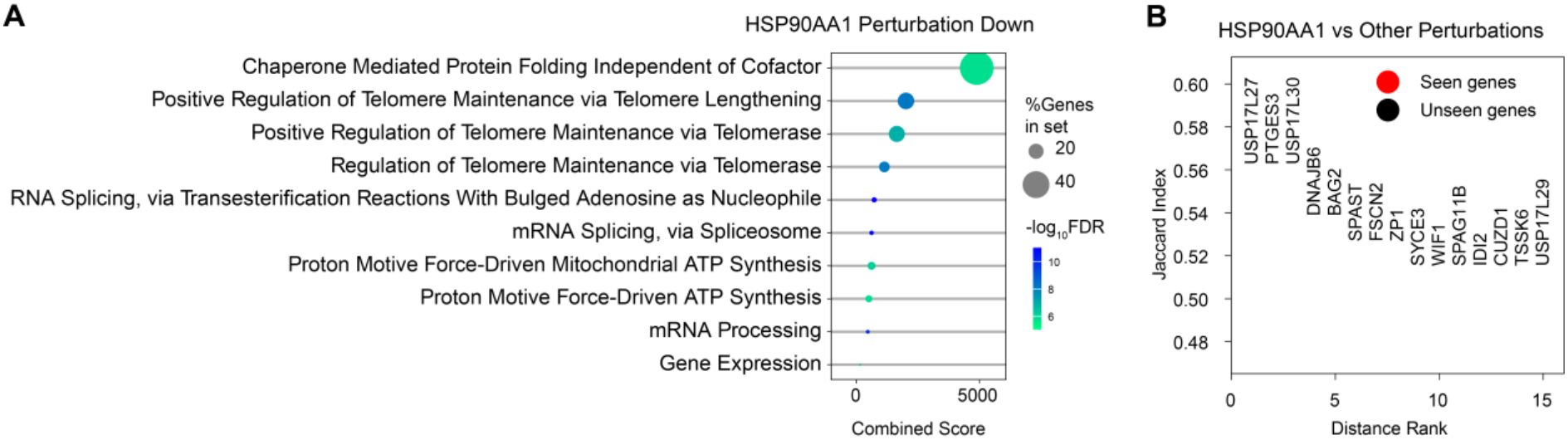
GFM predicts and prioritizes novel targets for HSP90AA1 knockdown in the K562 cell line. **(A)** Over-representation enrichment analysis of GO biological processes using GFM-predicted down-regulated genes with HSP90AA1 knockdown. The dot size shows the proportion of the genes in the enriched terms, and the color indicates log10-transformed 1/FDR. FDR: false discovery rate. **(B)** The predicted top 15 perturbations that are similar to HSP90AA1 knockdown based on the Jaccard index calculated with 100 down-regulated genes. The perturbations in red indicate the condition in the training set.

Perturbations were ranked according to the similarity of their predicted downregulated gene signatures to HSP90AA1 knockdown. Among the 15 highest-ranking perturbations were PTGES3, a canonical HSP90 co-chaperone, DNAJB6, a member of the HSP40 family, and BAG2, a regulator of chaperone activity. Notably, these predicted knockdown targets were also unseen during training but were predicted to induce similar transcriptional effects to HSP90AA1 knockdown (Fig. 5B). Taken together, these results indicate that GFM generalizes beyond individual perturbations to recover biologically accurate and functionally meaningful relationships suitable for *in silico* perturbation experiments.

## Discussion

In this study, we presented GFM, a biologically informed generative model for simulating post-perturbation single-cell transcriptomes. Building on previous work^10,11^, we incorporated GO, PPI, and perturbation similarity graphs into the model, enabling perturbation information to propagate and aggregate through biologically meaningful relationships. Unlike transformer-based perturbation embeddings, which rely on large-scale pretraining data to learn gene and perturbation relationships^7,8,28^, GFM directly incorporates biological priors through graph structures, providing inductive biases that facilitate learning in settings with limited perturbation data while allowing dataset- and context-specific optimization of perturbation representations. Beyond improving predictive performance, these biological graphs provide a structured framework for modeling how perturbations propagate through cellular systems. The perturbation similarity graph enables information sharing between perturbations with related transcriptional consequences, whereas GO and PPI graphs constrain gene-level information aggregation based on functional and physical relationships. It will be interesting to test additional biological priors as they become available, such as transcription factor regulatory networks, chromatin accessibility profiles, or ligand-receptor interaction networks. However, since GFM performance depends on perturbation-relevant graph connectivity (Fig. 4C, Supplementary Fig. 9), the utility of each prior will need to be evaluated for specific use cases rather than assuming that any additional prior knowledge improves prediction.

Our analyses show that the improved performance of GFM arises from the combination of complementary architectural components. First, GNNs introduce biologically informed inductive biases that facilitate information transfer between related genes and perturbations, reducing the dependence on large-scale perturbation datasets. Second, GFM leverages the latent flow matching framework^20^, which models cellular state generation by transforming a simple Gaussian source distribution into the target perturbed transcriptomic distribution through a deterministic ODE. Unlike approaches that construct flow trajectories between unperturbed and perturbed cells^8^, our model does not rely on explicit coupling between pre- and post-perturbation cells. Because scRNA sequencing is destructive, such one-to-one correspondences are fundamentally unobservable and must be approximated computationally, potentially introducing errors into the learned trajectories. Additionally, OT is used to couple Gaussian noise with perturbed latent representations on a per-condition basis before training, encouraging straighter flow trajectories while avoiding repeated expensive minibatch OT computations^13^ required during training in CellFlow^8^. Taken together, the integration of biologically informed graph representations with latent flow matching enables GFM to efficiently generate accurate single-cell perturbation responses while generalizing to unseen perturbations.

An effective latent flow matching model relies on a smooth and well-structured latent space, defined by a VAE in GFM. Designing an autoencoder that simultaneously captures generalizable biological variation and accurately reconstructs gene expression profiles remains challenging. Recent efforts have focused on developing universal single-cell foundation models trained with both observational and perturbational data^7,28–32^. However, their efficacy in predicting single-cell transcriptomes of out-of-distribution perturbations still requires further systematic evaluation. Here, we implemented a standard VAE to demonstrate the effectiveness of the GFM framework and provide a foundation for future integration of more advanced encoder architectures. We speculate that as improved representations of cellular states become increasingly more available, they will further enhance the ability of flow-based generative models to capture complex perturbation responses more accurately.

Notably, GFM consistently outperformed the simple yet competitive linear baseline, which itself outperforms all other current deep-learning models in predicting unseen single-gene perturbations, which is a fundamental phenomenon in the field^22^. We showed that GFM does not merely recapitulate generic perturbation-associated expression changes by comparing predictions against randomly selected perturbed cells from the training set^24^ (Fig. 2B, Fig. S1). In the two-gene perturbation prediction task, when none or one of the two tested genes was present in the training set (seen-0 and seen-1 subgroups), GFM outperformed the linear baseline and the other deep-learning models.

However, when both individual genes were observed during training (seen-2 subgroup), the additive model achieved near-perfect performance and consequently outperformed deep-learning-based approaches. This observation is consistent with the finding of Ahlmann-Eltze et al.^33^ that over 95% of the two-gene perturbations in the Norman dataset^3^ are additive, suggesting that models lacking explicit additive assumptions may be disadvantaged when training data are limited. As larger and more diverse perturbation atlases become available, frameworks such as GFM may become increasingly valuable for learning complex, non-additive cellular responses that cannot be captured by simple extrapolation from individual perturbations.

Collectively, our results demonstrate that accurate prediction of cellular responses to unseen perturbations requires integration of expressive generative models with biologically meaningful constraints. By combining knowledge graphs with latent flow matching, GFM provides a framework that captures both shared biological relationships among perturbations and the heterogeneity of single-cell transcriptomes. Particularly where datasets are limited, GFM can leverage existing biological knowledge to infer biological relationships, thereby improving generalization. As perturbation datasets continue to expand, biologically informed generative models may provide a scalable strategy for predicting cellular responses and prioritizing experimentally testable hypotheses in therapeutically relevant settings.

## Materials and Methods

### GFM

GFM is a deep learning framework designed to model and generate perturbation effects on single-cell transcriptomic profiles. GFM consists of three core modules: (i) a VAE to encode gene expression profiles into the latent space and decode them back to the gene expression space, (ii) a GNN for learning perturbation representations informed by biological knowledge graphs, and (iii) a neural ODE module to generate perturbed cells in the latent space conditioned by the GNN-derived perturbation representation.

Let *x* ∈ ℝ^*G*^ denote a cell-level gene expression vector with *G* genes, and let *y* denote the perturbation condition, which may contain one or more perturbed genes.

### VAE

We first map high-dimensional expression profiles into a low-dimensional latent space. The encoder parameterizes a multivariate Gaussian posterior

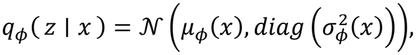

where *z* ∈ ℝ^*d*^ is a latent representation and *d* is the latent dimension. Latent samples are obtained with the reparameterization trick during training,

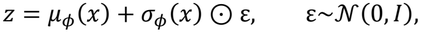

where *I* is an identity matrix. A decoder *f*_*θ*_ reconstructs the expression profile,

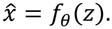

The VAE is trained by minimizing the evidence lower bound (ELBO):

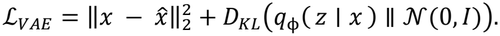

Only the cells in the training set will be used to pretrain the VAE. After pretraining, the estimated mean of the posterior *µ*_*ϕ*_(*x*) is directly used as the deterministic latent encoding for downstream flow matching training.

### Knowledge graph-constrained GAT

To represent perturbation structure, GFM constructs a perturbation graph *G*_*P*_ = (*V, E*), which is a concatenation of the GO, the PPI, and the perturbation similarity graphs. Within each graph, nodes correspond to genes and edges encode prior biological relationships. Each perturbation node *i* ∈ *V* is initialized with a learnable embedding 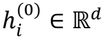. These node features are updated with 2 layers of GATs, enabling message passing and aggregation from 2-hop neighbor nodes.

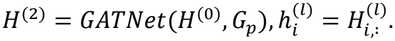

To represent a set of perturbations *S*, GFM aggregates node-level perturbation embeddings by summation:

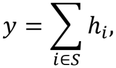

where *y* serves as the conditioning signal for the latent flow model. For the case of control cells, a vector of zeros is used as *y*.

### OT conditional flow matching

Given latent cell states and a perturbation condition, the neural ODE module learns a time-dependent velocity field that transports Gaussian noise to target perturbed states in the latent space. To enable efficient training and improve performance on conditional data, we leverage precomputed OT plans for each condition before model training^34,35^. Specifically, before optimizing the flow model, GFM computes exact OT couplings separately within each perturbation condition, yielding condition-specific paired samples (*z*_0_, *z*_1_) ∼ *π*^(*y*)^, where *z*_0_ denotes Gaussian noise and *z*_1_ denotes a target latent state. These precomputed couplings are then used throughout training. Conditioned on the perturbation embedding and time, the model learns a velocity field

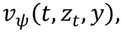

where *t* ∈ [0, 1], *z*_*t*_ is an intermediate latent state along the path, and *y* is the perturbation label. For a coupled pair (*z*_0_, *z*_1_) and time *t* ∼ *Uniform*(0, 1), GFM uses an affine interpolation path

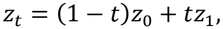

with target velocity

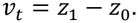

The flow model is trained by regressing the predicted velocity to the target velocity with the conditional flow matching (CFM) loss:

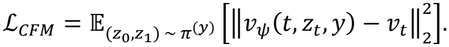

In the implementation, time is encoded with sinusoidal embeddings^36^, and the latent state, perturbation embedding, and the time embedding are concatenated before being passed to a MLP that outputs the velocity vector. The GNN module and the neural ODE module are trained jointly with the CFM loss.

During inference, we implement classifier-free guidance^37^ by combining conditional and unconditional velocity fields:

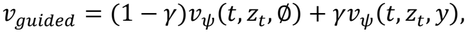

where we pick *γ* = 1.5 as the guidance scale and ∅ is a vector of zeros representing the unperturbed condition. GFM generates perturbed latent states by integrating *v*_*guided*_ from *t* = 0 to *t* = 1 using the Dormand-Prince (dopri5) ODE solver^38^. The final latent state *z*_1_ is then decoded by the pretrained VAE decoder to obtain the predicted gene expression profile:

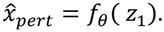

### Graph construction

GFM incorporates several biologically informed perturbation graphs. In all cases, nodes are genes, and each target node retains its top-k highest-weight incoming neighbors. GFM concatenates multiple biological knowledge graphs to form a joint graph *G*_*p*_.

### GO graph

The GO similarity graph is directly obtained from GEARS^10^. Each gene is represented by its annotated GO term set. Pairwise similarity between genes is computed with the Jaccard index,

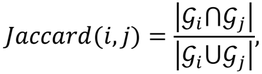

where *G*_*i*_ and *G*_*j*_ are the GO-term sets for genes *i* and *j*. For each target gene, GFM keeps the top 20 most similar source genes.

### PPI graph

For the protein-protein interaction (PPI) graph, edges are taken from STRING database (human, v12.0)^21^ with associated combined scores. For each target gene, the model retains the top 10 source genes with the largest combined score.

### Perturbation similarity graph

For the perturbation-similarity graph, GFM constructs a transcriptional signature for each perturbed gene from the Replogle K562 genome-wide perturbation dataset^4^. First, the mean expression profile is computed for each perturbation condition and for the control condition. The corresponding perturbation signature for perturbation *i* is the control-relative difference

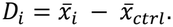

The pairwise similarity between perturbations is computed with cosine similarity,

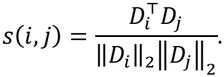

For each target gene, the top 20 similar perturbations are connected.

## Baseline models

### Linear model

We model each perturbation as an additive shift from the matched control expression state. We first aggregate single-cell expression profiles into means for each perturbation-condition. Let *x*_*p*_ denote the mean expression vector for perturbation *p* and let *x*_*ctrl*_ denote the control mean. We derive perturbation embeddings by applying PCA to the transpose of the mean expression matrix of the training data, producing a gene-level PCA embedding matrix *E* ∈ ℝ^*G*×*d*^ with *d* = 50 components. For perturbation *p*, a multi-hot vector *m*_*p*_ selects the perturbed gene or genes, and the perturbation embedding is computed as:

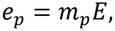

where *e*_*p*_ is the PCA-based perturbation embedding. A linear model was then trained to predict the perturbation-induced change in expression:

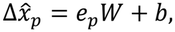

where *W* ∈ ℝ^*d*×*G*^ is the learned weight matrix, and *b* ∈ ℝ^*G*^ is the learned bias vector. The model was optimized by minimizing the MSE between the observed shift and the predicted shift:

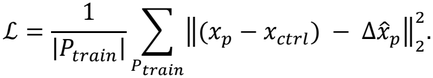

Finally, the predicted perturbed expression profile was obtained by adding the predicted perturbation effect to the matched control state:

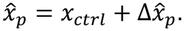

### Perturb train

To avoid biasing toward systemic variance in each dataset, we randomly select 100 perturbed cells from the training set for each perturbation as the output of the perturb train model^24^.

### Technical duplicates

To use empirical data as the positive control for the evaluation metrics, we randomly split the ground-truth cells for each perturbation into two halves and use one half as the prediction to calculate the evaluation metrics with the other half^33^.

### Additive model

In the “seen-2” subgroup of the test set in the two-gene perturbation prediction task, the output perturbation shift is the summation of each perturbation in the training set. Let 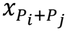 be the mean gene expression of perturbation *i* and *j*, and *x*_*ctrl*_ be the mean expression of control cells. The additive model output is

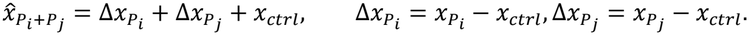

## Evaluation metrics

We select the top *k* DEGs for each perturbation to evaluate model performance. Let *X* ∈ ℝ^*m*×*k*^ denote the ground-truth expression matrix and *Y* ∈ ℝ^*n*×*k*^ denote the predicted expression matrix of the selected DEGs for a given perturbation.

### MSE

MSE is computed between the gene-wise mean expression vectors of the true and the predicted cells,

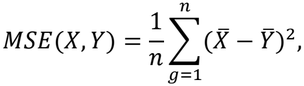

where 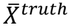 and 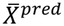 are the mean expression profiles over the cells in each perturbation.

### Pearson correlation coefficient (*r*)

Pearson correlation coefficient is computed between these same mean expression vectors,

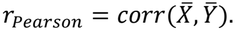

### Pearson delta

To quantify whether the predicted perturbation effect matched the true shift, we also compute Pearson delta correlation. Specifically, we use the mean expression profile from randomly selected perturbed cells in the training set (perturb-train) 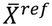 as the reference. We first subtracted 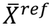 from both the true and predicted mean profiles, and then computed Pearson correlation on the resulting differential signals,

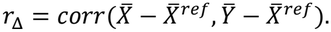

This metric emphasizes agreement in perturbation-induced changes rather than absolute expression levels.

### 2-WD

To evaluate distributional similarity, we first use the 2-WD to measure the cheapest transport cost of moving the true cell distribution to the predicted one. The implementation constructs the squared Euclidean cost matrix

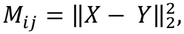

assumes uniform weights over cells in each perturbation, and computes the OT cost

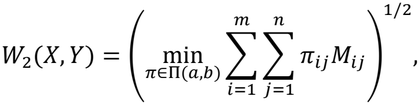

where *a*_*i*_ = 1/*m, b*_*j*_ = 1/*n*, and Π(*a, b*) denotes the set of all transport plans with marginals *a* and *b*.

### E-distance

Finally, we compute the E-distance that compares the pairwise Euclidean distances between two distributions and within each distribution, which not only measures the distance between two distributions but also accounts for the variations within them^39^. Formally, the pairwise Euclidean distances are defined as

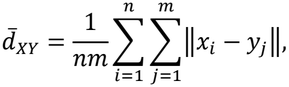

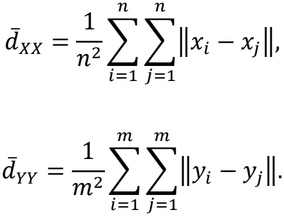

The E-distance 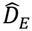 is

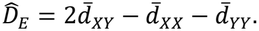

## Data preprocessing

For the Perturb-seq data from Replogle et al.^4^ (K562, RPE1, K562 genome-wide perturbation screening), we followed GEARS’ approach to filter for strong perturbations. Specifically, perturbations were retained if they exhibited more than 50 differentially expressed genes, a percent knockdown of at least 30%, and at least 25 cells. We further removed individual cells that did not show evidence of knockdown at the single-cell level. For each perturbed gene, only cells in which the expression of the targeted gene was at or below the 10th percentile of its expression in non-targeting control cells were retained. HepG2 and Jurkat datasets from Nadig et al.^5^ were pre-filtered in STATE^31^ for strong knockdown efficiency. For the two-gene perturbation dataset from Norman et al.^3^, we followed the steps in GEARS ([GEARS Norman preprocessing](https://github.com/yhr91/GEARS_misc/blob/f88211870dfa89c38a2eedbd69ca1abd28a25f3c/data/preprocessing/Norman19.ipynb)) to reformat the metadata without further filtering.

Following perturbation-level filtering, all perturb-seq data were normalized by library size and log1p transformed. The top 5000 highly variable genes and all perturbed genes were retained for downstream analysis. Per-condition differentially expressed genes relative to the non-targeting control were ranked by absolute effect size using a Wilcoxon rank-sum test^40^.

## Training and prediction setup

For each single-gene perturbation dataset, perturbations were split into train (75%), validation (10%), and test (15%) sets (Supplementary Table 2). For the Norman two-gene perturbation dataset^3^, ‘*prepare_split*’ function with ‘*split=’simulation‘*’) from ‘*gears*’ was used to create the data splits, ending up with 50% of the perturbations in the training, 10% in the validation, and 40% in the test sets (Supplementary Table 3). The perturbations in the test set were held out during training and used only for evaluating model performance. The validation set helped prevent the model from overfitting to the training set. Three random splits for each dataset were used across all models. To ensure fair comparisons, the number of predicted cells from all models was set to one hundred for each perturbation. Each experiment was run on 1 GPU slice with 18 GB memory from 4 × NVIDIA H200 NVL GPUs (driver 610.43.02, CUDA 13.3).

### GFM

GFM training consists of two stages: pretraining of the VAE and training of the latent flow matching module. For VAE pretraining, only cells in the training set are used for optimization. Training runs for a maximum of 500 epochs and is stopped early if the validation loss does not decrease for 100 consecutive epochs. The resulting latent cell representations are then used for OT coupling computation and flow matching training as described above. The flow matching model is trained for a maximum of 500 epochs. Every 50 epochs, the predicted latent cell representations of the validation set are evaluated using the 2-WD. Training is stopped early if the validation 2-WD does not decrease over four consecutive evaluations.

### CellFlow (cellflow-tools v0.0.7)

For CellFlow implementation, ESM2 embeddings for all perturbed genes were first generated following the author’s instructions ([CellFlow ESM embedding](https://github.com/theislab/CellFlow/blob/docs/tutorial/satija/docs/notebooks/300_satija_unseen_cell_line_cytokine.ipynb)). Default settings from the CellFlow tutorial ([CellFlow GitHub](https://github.com/theislab/CellFlow/tree/main)) were then applied for model training and unseen perturbation prediction.

### GEARS (cell-gears v0.1.2)

We followed the GEARS tutorial ([GEARS GitHub](https://github.com/snap-stanford/GEARS)) for model training. To obtain gene expression predictions at the single-cell level, we removed the final averaging step from the original prediction function.

### scGPT (scgpt v0.2.4)

We followed the scGPT tutorial ([Fine-tuning Pre-trained Model for Perturbation Prediction — scGPT 0.2.1 documentation](https://scgpt.readthedocs.io/en/latest/tutorial_perturbation.html)) to fine-tune the pretrained model for prediction. We downloaded the whole-human pretrained model from bowang-lab/scGPT. The final averaging step in the original prediction function was also removed to obtain single-cell gene expression predictions.

## Ablation experiments

For the ablation of GFM modules, we examined the role of the flow-matching objective and graph topology. To disable flow matching, we replaced the OT velocity-matching objective with direct regression from the source latent state *z*_0_ to the target latent state *z*_1_. Specifically, the neural ODE module took uncoupled Gaussian noise and a sinusoidal time embedding^36^ at *t* = 0 as the inputs and was optimized with an MSE loss between the predicted latent endpoint and the target latent state.

This preserves the same conditional architecture but removes the OT coupling and the continuous transport formulation. To remove biologically informed graph topology, we kept the same number of nodes and edges but randomly permuted graph source nodes, thereby disrupting biologically meaningful connectivity while preserving graph density and degree scale. We performed both operations if both modules were ablated.

For the ablation of different types of knowledge graphs, we evaluated the contribution of each biological graph while keeping the rest of the model fixed. As mentioned above, we assessed whether predictive performance is driven primarily by GO, PPI, perturbation similarity, or their combination by randomizing the selected biological graph(s).

For all ablation experiments, model performance was assessed on held-out test conditions.

## Graph connectivity analysis

We quantified whether local graph connectivity was associated with predictive performance in GFM. For each held-out test condition, we extracted upstream source nodes by traversing incoming edges in the directed graph for two hops, corresponding to the local neighborhood used in this analysis. We then identified which source nodes had been observed in the training split and recorded two connectivity statistics: the number of trained source nodes and the fraction of all source nodes that were present in training. To quantify transcriptional similarity between a test perturbation and its trained graph neighbors, we computed the mean expression profile for each condition, subtracted the mean control profile to obtain a perturbation effect vector, and calculated cosine similarity between the held-out condition and each trained source-node perturbation reference. For each test condition, we summarized these similarities by their maximum and median values. This analysis was performed using three predefined data splits of the indicated single-gene perturbation datasets.

## CML target identification

GFM was trained on the Replogle K562 genome-wide perturbation screen dataset^4^ using a 90% training and 10% validation split and no held-out test set. The GO graph, PPI graph, and the perturbation similarity graph as mentioned above were used here. The predictions were generated for every perturbation node in the graph. For each perturbation, 20 synthetic single-cell profiles were sampled for downstream analysis. Predicted cells were then aggregated to perturbation-level mean expression profiles. The resulting perturbation means were embedded by PCA followed by Leiden clustering and t-SNE visualization. Cluster identities were annotated by ranking differentially expressed genes across Leiden clusters with a Wilcoxon test and performing Gene Ontology Biological Process^41^ enrichment on the 100 most downregulated genes per cluster.

For target identification, predicted perturbation profiles were compared against sampled control cells by differential expression analysis. Using HSP90AA1 as the query perturbation, we quantified similarity between the query signature and every other perturbation by the Jaccard index computed on the two sets of 100 downregulated genes. Functional interpretation of the query signature was obtained by Gene Ontology enrichment analysis of the same 100 downregulated genes using ‘gseapy’^42^.

## Data availability

Norman^3^, Replogle K562, Replogle RPE1, Replogle K562 genome-wide perturbation screen^4^ datasets were obtained via pertpy package^43^ (v0.10.0). The pre-filtered Nadig HepG2 and Nadig Jurkat datasets^5^ were downloaded from Arc Institute Hugging Face repository Replogle-Nadig-Preprint (arcinstitute/Replogle-Nadig-Preprint at main). The GO graph was obtained from GEARS^10^ (https://dataverse.harvard.edu/api/access/datafile/6934319). The human PPI data for graph construction was downloaded from STRING database v12.0^21^ (string-db.org).

## Code availability

The GFM package and the complete analysis scripts are available via GitHub at TRON-Bioinformatics/GFM. GFM is distributed under the MIT License.

## Acknowledgements

This work was supported by the German Research Council (DFG) (496710327) and by the German Federal Ministry of Research, Technology and Space (BMFTR) within the framework of the project curATime (03ZU2202DA, 03ZU2202GB, 03ZU1202JA).

## Author information

## Contributions

H.L. conceptualized the study, developed GFM, performed analysis, and wrote the manuscript. B.Sch., P.R.F, J.V.G., M.S., D.A.S, G.G., S.V., B.St., and J.K. provided overall input and guidance. J.K. and B.St. supervised the project, and all authors contributed to writing the paper.

## References

1. Adamson, B. et al. A Multiplexed Single-Cell CRISPR Screening Platform Enables Systematic Dissection of the Unfolded Protein Response. Cell 167, 1867–1882.e21 (2016).

2. Dixit, A. et al. Perturb-Seq: Dissecting Molecular Circuits with Scalable Single-Cell RNA Profiling of Pooled Genetic Screens. Cell 167, 1853–1866.e17 (2016).

3. Norman, T. M. et al. Exploring genetic interaction manifolds constructed from rich single-cell phenotypes. Science 365, 786–793 (2019).

4. Replogle, J. M. et al. Mapping information-rich genotype-phenotype landscapes with genome-scale Perturb-seq. Cell 185, 2559–2575.e28 (2022).

5. Nadig, A. et al. Transcriptome-wide analysis of differential expression in perturbation atlases. Nat. Genet. 57, 1228–1237 (2025).

6. Gavriilidis, G. I., Vasileiou, V., Orfanou, A., Ishaque, N. & Psomopoulos, F. A mini-review on perturbation modelling across single-cell omic modalities. Comput. Struct. Biotechnol. J. 23, 1886–1896 (2024).

7. Cui, H. et al. scGPT: toward building a foundation model for single-cell multi-omics using generative AI. Nat. Methods 21, 1470–1480 (2024).

8. Klein, D. et al. CellFlow enables generative single-cell phenotype modeling with flow matching. Preprint at 10.1101/2025.04.11.648220 (2025).

9. Lin, Z. et al. Evolutionary-scale prediction of atomic-level protein structure with a language model. Science 379, 1123–1130 (2023).

10. Roohani, Y., Huang, K. & Leskovec, J. Predicting transcriptional outcomes of novel multigene perturbations with GEARS. Nat. Biotechnol. 42, 927–935 (2024).

11. Wenkel, F. et al. TxPert: using multiple knowledge graphs for prediction of transcriptomic perturbation effects. Nat. Biotechnol. https://doi.org/10.1038/s41587-026-03113-4 (2026) doi:10.1038/s41587-026-03113-4.

12. Lipman, Y., Chen, R. T. Q., Ben-Hamu, H., Nickel, M. & Le, M. Flow Matching for Generative Modeling. Preprint at 10.48550/ARXIV.2210.02747 (2022).

13. Tong, A. et al. Improving and generalizing flow-based generative models with minibatch optimal transport. Preprint at 10.48550/ARXIV.2302.00482 (2023).

14. Rohbeck, M. et al. Modeling complex system dynamics with flow matching across time and conditions. in International Conference on Learning Representations vol. 2025 12164–12188 (2025).

15. Wang, D. et al. Joint Velocity-Growth Flow Matching for Single-Cell Dynamics Modeling. In The Thirty-ninth Annual Conference on Neural Information Processing Systems (2025).

16. Atanackovic, L. et al. Meta Flow Matching: Integrating Vector Fields on the Wasserstein Manifold. in The Thirteenth International Conference on Learning Representations (2025).

17. Rubbi, A. et al. MixFlow: Mixture-Conditioned Flow Matching for Out-of-Distribution Generalization. CoRR abs/2601.11827, (2026).

18. Rombach, R., Blattmann, A., Lorenz, D., Esser, P. & Ommer, B. High-Resolution Image Synthesis with Latent Diffusion Models. Preprint at 10.48550/ARXIV.2112.10752 (2021).

19. Labs, B. F. FLUX.2: Frontier Visual Intelligence. (2025).

20. Dao, Q., Phung, H., Nguyen, B. & Tran, A. Flow Matching in Latent Space. Preprint at 10.48550/arXiv.2307.08698 (2023).

21. Szklarczyk, D. et al. The STRING database in 2023: protein–protein association networks and functional enrichment analyses for any sequenced genome of interest. Nucleic Acids Res. 51, D638–D646 (2023).

22. Ahlmann-Eltze, C., Huber, W. & Anders, S. Deep-learning-based gene perturbation effect prediction does not yet outperform simple linear baselines. Nat. Methods 22, 1657–1661 (2025).

23. Miller, H. E. et al. Deep learning perturbation models can outperform baselines on calibrated metrics. Nat. Biotechnol. https://doi.org/10.1038/s41587-026-03307-w (2026) doi:10.1038/s41587-026-03307-w.

24. Viñas Torné, R. et al. Systema: a framework for evaluating genetic perturbation response prediction beyond systematic variation. Nat. Biotechnol. https://doi.org/10.1038/s41587-025-02777-8 (2025) doi:10.1038/s41587-025-02777-8.

25. Veličković, P. et al. Graph Attention Networks. Preprint at 10.48550/ARXIV.1710.10903 (2017).

26. Bhatia, S. et al. Targeting HSP90 dimerization via the C terminus is effective in imatinib-resistant CML and lacks the heat shock response. Blood 132, 307–320 (2018).

27. Holt, S. E. et al. Functional requirement of p23 and Hsp90 in telomerase complexes. Genes Dev. 13, 817–826 (1999).

28. Szałata, A. et al. PerturBERT: Learning Gene Co-Variation Embeddings from Perturbation Signatures. in ICLR 2026 Workshop on Machine Learning for Genomics Explorations (2026).

29. Palla, G. et al. Scalable Single-Cell Gene Expression Generation with Latent Diffusion Models. Preprint at 10.48550/ARXIV.2511.02986 (2025).

30. Dong, M. et al. Stack: In-Context Learning of Single-Cell Biology. Preprint at 10.64898/2026.01.09.698608 (2026).

31. Adduri, A. K. et al. Predicting cellular responses to perturbation across diverse contexts with State. Preprint at 10.1101/2025.06.26.661135 (2025).

32. Rosen, Y. et al. Universal cell embedding provides a foundation model for cell biology. Nature https://doi.org/10.1038/s41586-026-10689-z (2026) doi:10.1038/s41586-026-10689-z.

33. Miller, H. E. et al. Deep Learning-Based Genetic Perturbation Models Do Outperform Uninformative Baselines on Well-Calibrated Metrics. Preprint at 10.1101/2025.10.20.683304 (2025).

34. Cheng, H. K. & Schwing, A. The curse of conditions: Analyzing and improving optimal transport for conditional flow-based generation. in Proceedings of the IEEE/CVF International Conference on Computer Vision 15875–15884 (2025).

35. Hui, K.-H., Liu, C., Fu, C.-W., Vahdat, A., & others. Not-so-optimal transport flows for 3d point cloud generation. in International Conference on Learning Representations vol. 2025 98748–98771 (2025).

36. Lipman, Y. et al. Flow Matching Guide and Code. Preprint at 10.48550/ARXIV.2412.06264 (2024).

37. Zheng, Q. et al. Guided Flows for Generative Modeling and Decision Making. (2023).

38. Chen, R. T. Q. torchdiffeq. (2018).

39. Peidli, S. et al. scPerturb: harmonized single-cell perturbation data. Nat. Methods 21, 531–540 (2024).

40. Wolf, F. A., Angerer, P. & Theis, F. J. SCANPY: large-scale single-cell gene expression data analysis. Genome Biol. 19, 15 (2018).

41. The Gene Ontology Consortium et al. The Gene Ontology knowledgebase in 2026. Nucleic Acids Res. 54, D1779–D1792 (2026).

42. Fang, Z., Liu, X. & Peltz, G. GSEApy: a comprehensive package for performing gene set enrichment analysis in Python. Bioinformatics 39, btac757 (2023).

43. Heumos, L. et al. Pertpy: an end-to-end framework for perturbation analysis. Nat. Methods 23, 350–359 (2026).

