## Supplementary information for "Graph-guided flow matching enables biologically informed prediction at single-cell resolution"

### Supplementary figures

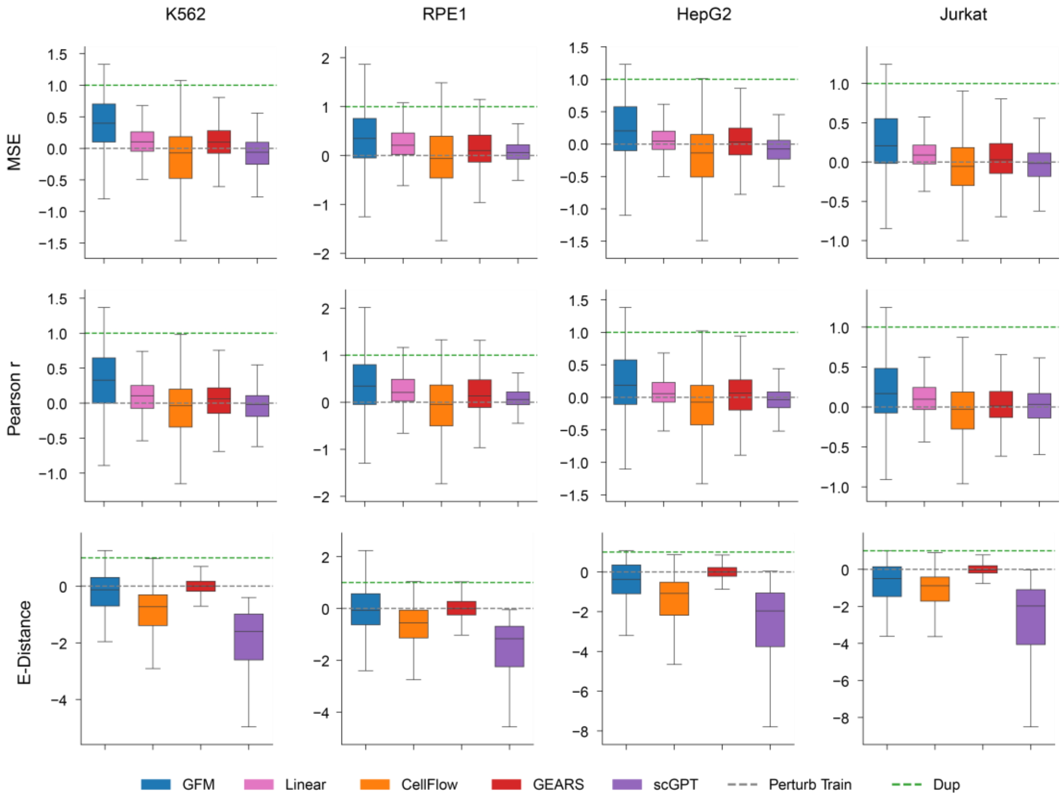


**Supplementary Fig. 1.** Box plots showing MSE, Pearson r, and E-distance of the predicted top 20 DEGs of each perturbation for each model in K562, RPE1, HepG2, and Jurkat datasets. Randomly selected perturbed cells from the training set (Perturb train) and technical duplicate (Dup) were used to normalize the metrics. The tested perturbations in three independent data splits were pooled.

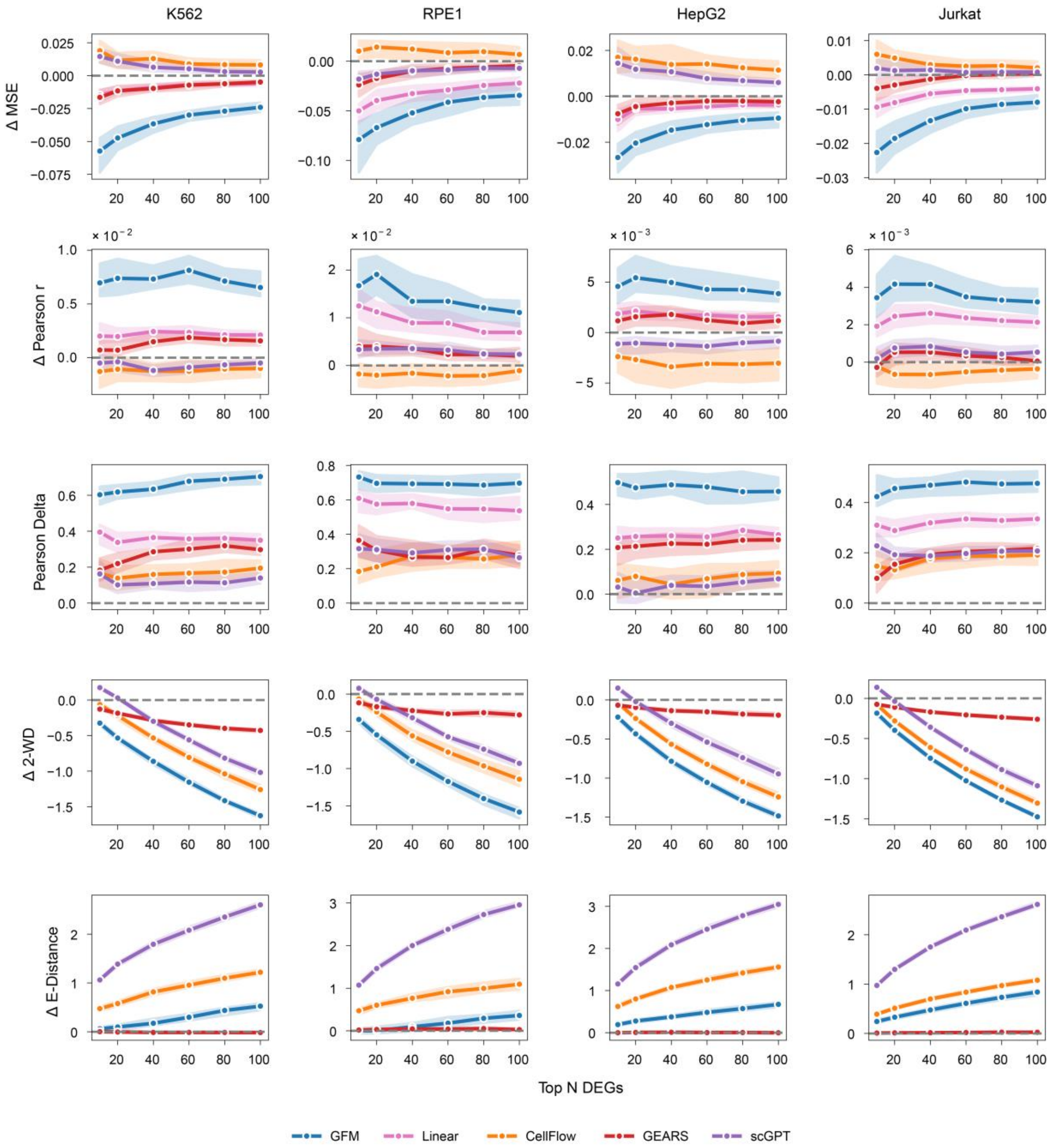


**Supplementary Fig. 2.** Line plots of delta MSE, delta Pearson r, Pearson delta, delta 2-WD, and delta E-distance of the predicted top 10, 20, 40, 60, 80, and 100 DEGs of each perturbation relative to Perturb train baseline. The analysis was performed in the same way for each model in the K562, RPE1, HepG2, and Jurkat datasets. Results calculated from three independent data splits were pooled, and the median and 95% confidence interval were shown.

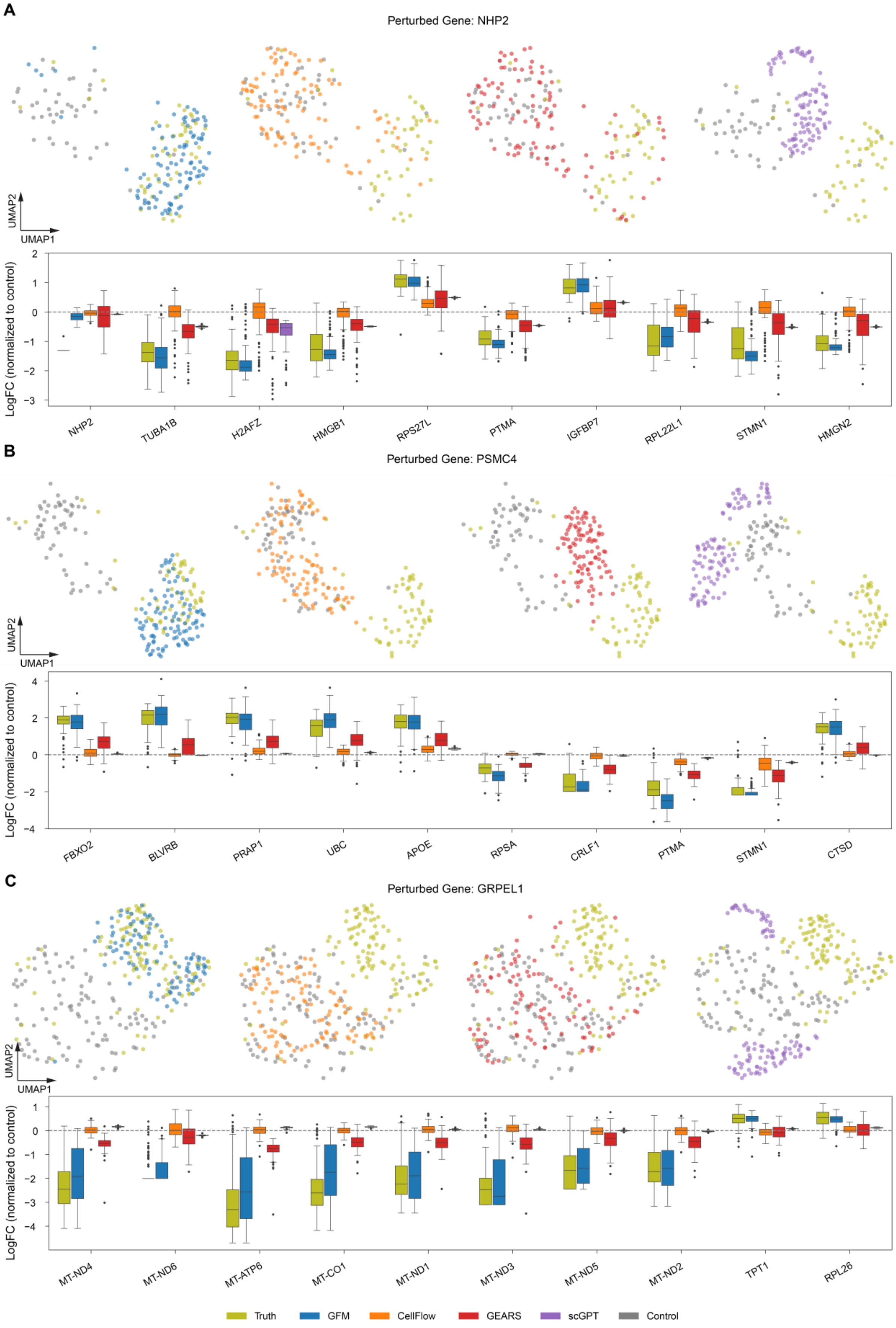
A
Perturbed Gene: NHP2
UMAP2
UMAP1
LogFC (normalized to control)
NHP2
TUBA1B
H2AFZ
HMGB1
RPS27L
PTMA
IGFBP7
RPL22L1
STMN1
HMGN2
B
Perturbed Gene: PSMC4
UMAP2
UMAP1
LogFC (normalized to control)
FBXO2
BLVRB
PRAP1
UBC
APOE
RPSA
CRLF1
PTMA
STMN1
CTSD
C
Perturbed Gene: GRPEL1
UMAP2
UMAP1
LogFC (normalized to control)
MT-ND4
MT-ND6
MT-ATP6
MT-CO1
MT-ND1
MT-ND3
MT-ND5
MT-ND2
TPT1
RPL26
Truth
GFM
CellFlow
GEARS
scGPT
Control

**Supplementary Fig. 3.** Joint UMAP representations and the predicted top 10 DEGs of NHP2 knockdown in RPE1 (**A**), PSMC4 knockdown in HepG2 (**B**), and GRPEL1 knockdown in Jurkat (**C**) from different models. Expression levels were shown as log fold change relative to the mean control cell expression. Truth: ground-truth gene expression.

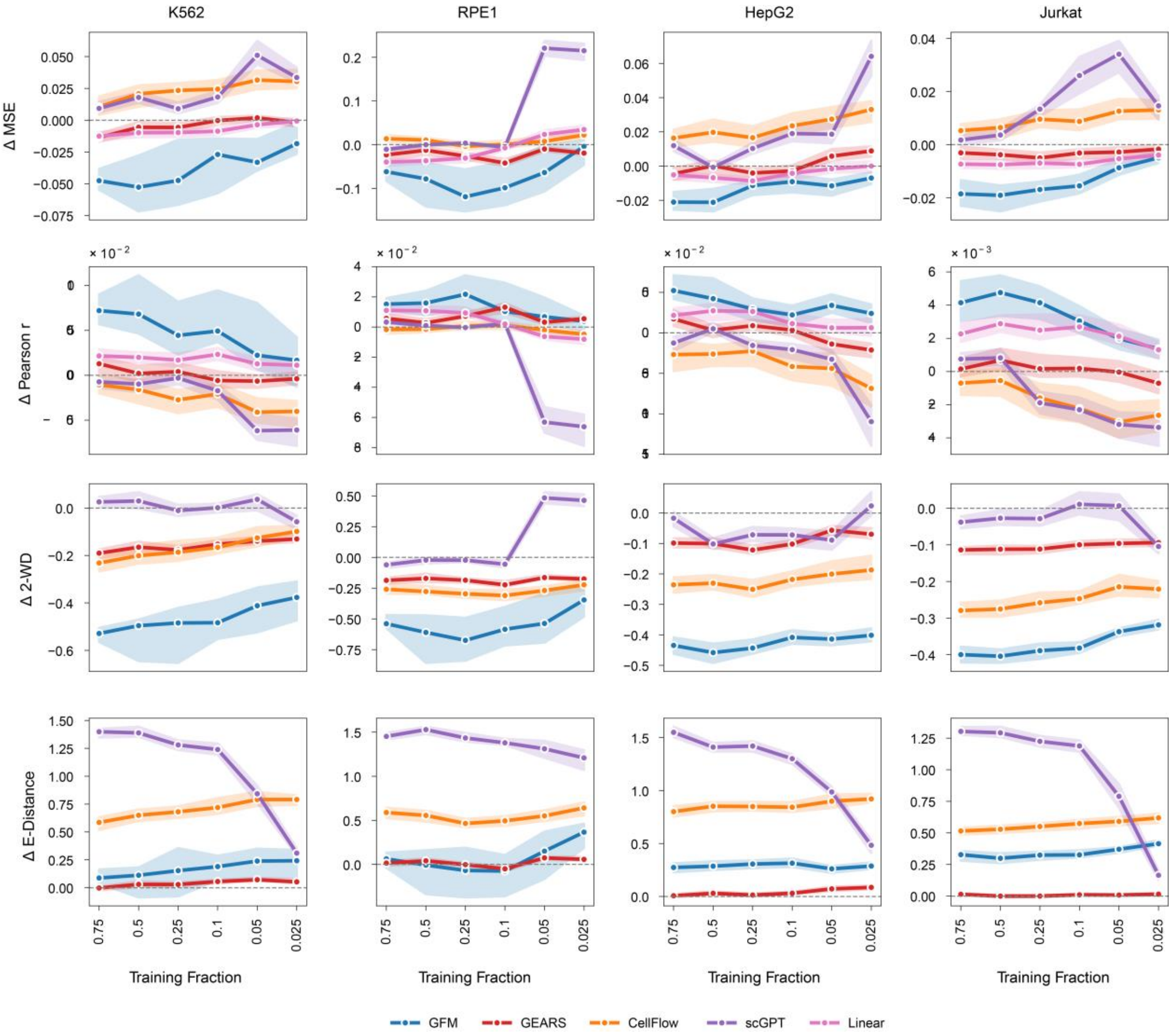


**Supplementary Fig. 4.** Line plots of delta MSE, delta Pearson r, delta 2-WD, and delta E-distance of the predicted top 20 DEGs of each perturbation with reduced training fractions (0.75, 0.5, 0.25, 0.1, 0.05, and 0.025) relative to Perturb train baseline. The analysis was performed in the same way for each model in the K562, RPE1, HepG2, and Jurkat datasets. Results calculated from three independent data splits were pooled, and the median and 95% confidence interval were shown.

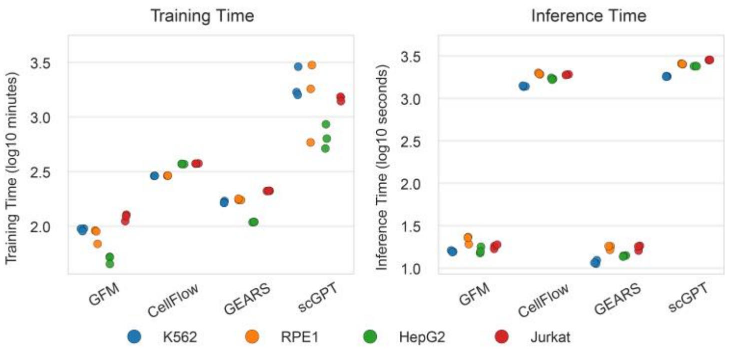


**Supplementary Fig. 5.** Computation time of training and inference of each model using K562, RPE1, HepG2, and Jurkat datasets with three data splits.

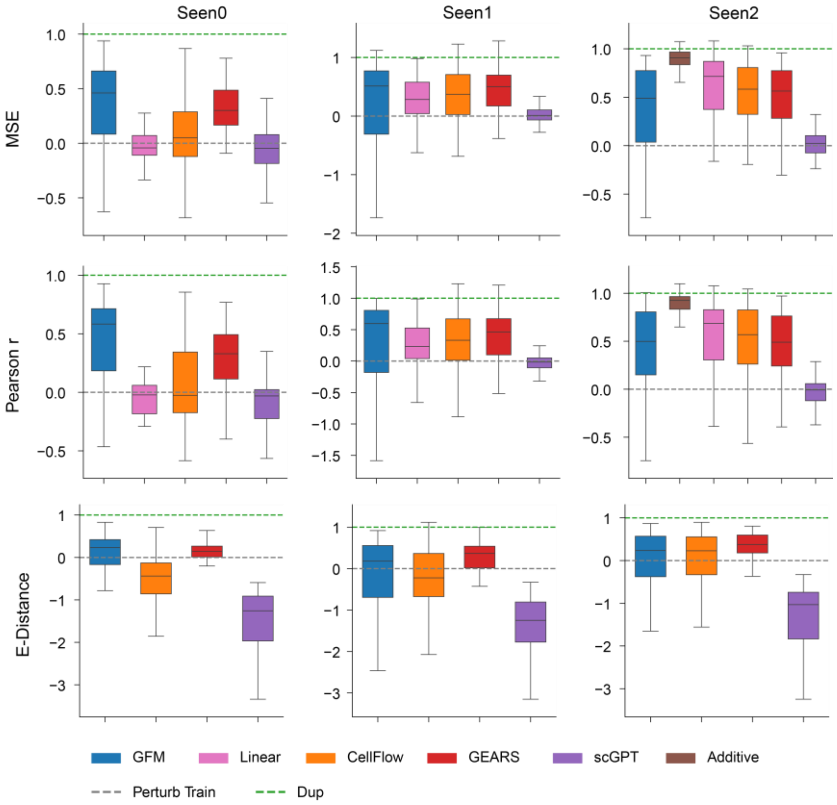


**Supplementary Fig. 6.** Box plots showing MSE, Pearson r, an E-distance of the predicted top 20 DEGs of each perturbation for each model in seen-0, seen-1, and seen-2 subgroups. Randomly selected perturbed cells from the training set (Perturb train) and technical duplicate (Dup) were used to normalize the metrics. The tested perturbations in three independent data splits were pooled.

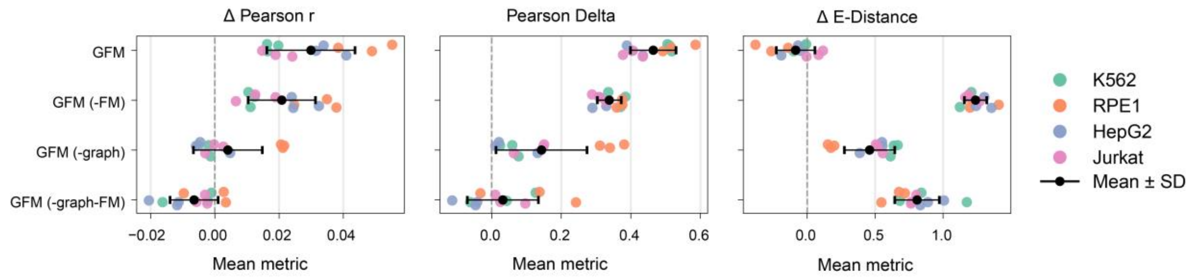


**Supplementary Fig. 7.** Additional metrics for GFM module ablation studies. The aggregated mean delta Pearson r, Pearson delta, and delta E-distance relative to Perturb train baseline for the three data splits in K562, RPE1, HepG2, and Jurkat datasets in different ablation conditions. Additional means and standard deviations were shown for each ablation condition. -FM: without OT flow matching, -graph: randomly permuted graph.

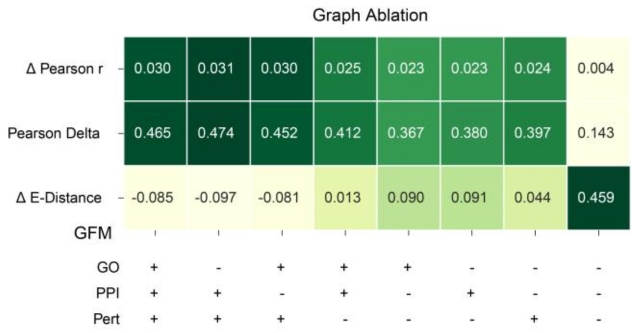


**Supplementary Fig. 8.** A heatmap showing the averaged delta Pearson r, Pearson delta, and delta E-distance across relative to Perturb train baseline three independent splits of K562, RPE1, HepG2, and Jurkat datasets under multiple graph ablation conditions. The indicated biological graph was randomly permuted (-) or remained intact (+).

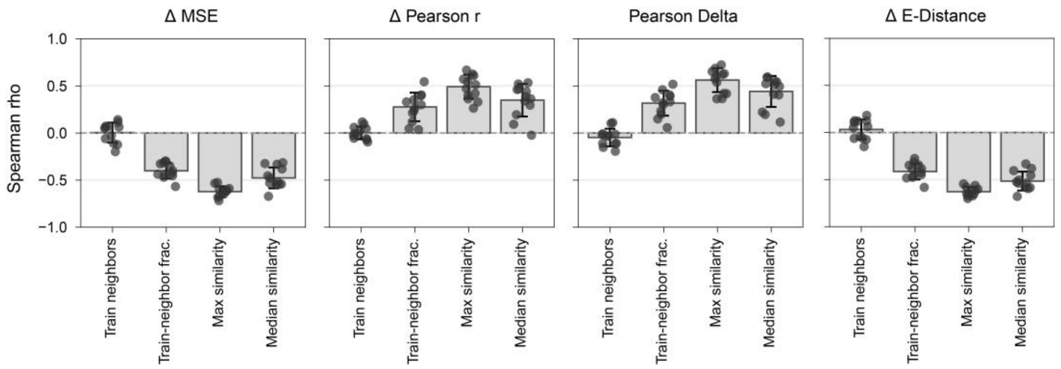


**Supplementary Fig. 9.** Summarized Spearman correlation between the graph-derived properties and metrics, including delta MSE, delta Pearson r, Pearson delta, and delta E-distance. Each data point shows the Spearman rho of a data split in the K562, RPE1, HepG2, and Jurkat datasets. For each comparison group, the Fisher z-transformed mean of the Spearman rho was shown as the bar, with the error bar showing the standard deviation.

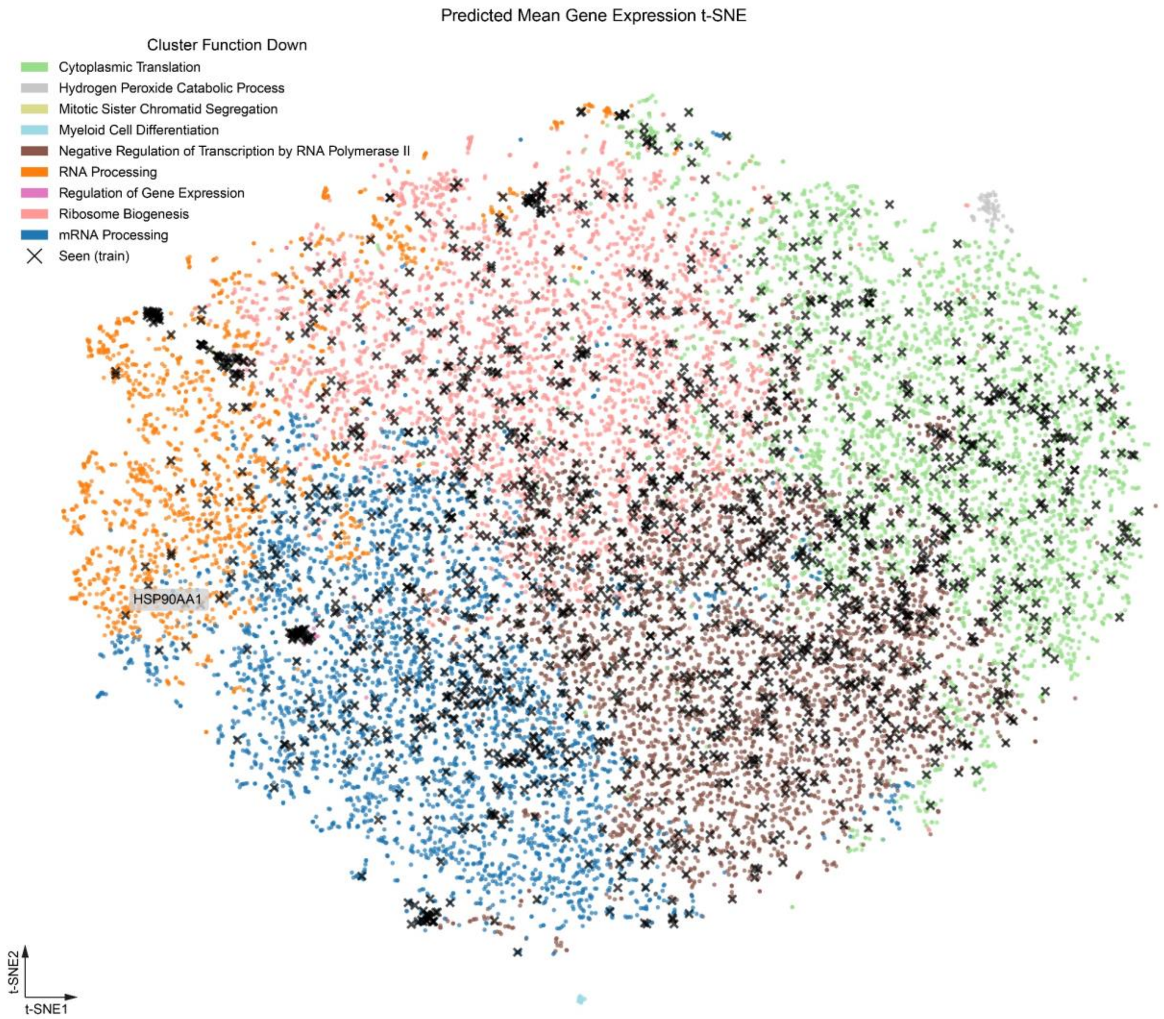


**Supplementary Fig. 10.** A t-SNE representation of the GFM prediction of the entire perturbation space for K562. Each data point represents the mean predicted gene expression of a perturbation and was further clustered and annotated with Gene Ontology Biological Process terms by the top 100 predicted down-regulated genes. Perturbations present in the training split were labeled with crosses.

### Supplementary tables

**Supplementary Table 1.** Number of conditions in the training set of each single-gene perturbation dataset for the experiment of reducing training size (including control).

| Percentage of training set | Replogle K562 | Replogle RPE1 | Nadig HepG2 | Nadig Jurkat |
|---|---|---|---|---|
| 75% | 820 | 1158 | 1006 | 1154 |
| 50% | 547 | 773 | 671 | 770 |
| 25% | 274 | 387 | 336 | 385 |
| 10% | 110 | 155 | 135 | 154 |
| 5% | 55 | 78 | 68 | 77 |
| 2.5% | 88 | 39 | 34 | 39 |

**Supplementary Table 2.** Number of conditions in the train, validation, and test sets of one-gene perturbation datasets (including control in the training set).

| Partition | Replogle K562 | Replogle RPE1 | Nadig HepG2 | Nadig Jurkat |
|---|---|---|---|---|
| Train (75%) | 820 | 1158 | 1006 | 1154 |
| Validation (10%) | 109 | 154 | 134 | 154 |
| Test (15%) | 164 | 232 | 201 | 230 |
| Total (100%) | 1093 | 1544 | 1341 | 1538 |

**Supplementary Table 3.** Number of conditions in each data split for the Norman two-gene perturbation dataset, including subgroups in the test set (including control in the training set).

| Partition | Split 1 | Split 2 | Split 3 |
|---|---|---|---|
| Train (~50%) | 136 | 140 | 138 |
| Validation (~10%) | 34 | 20 | 33 |
| Test (~40%) | 110 | 120 | 109 |
| Seen 0 | 9 | 12 | 6 |
| Seen 1 | 44 | 53 | 49 |
| Seen 2 | 20 | 17 | 19 |
| Unseen single | 37 | 38 | 35 |
